# Comparative Analysis of 3D Culture Methodologies in Prostate Cancer Cells

**DOI:** 10.1101/2025.03.11.642589

**Authors:** Ella Foster, Oliver Wardhana, Ziyu Zeng, Xin Lu

## Abstract

Three-dimensional (3D) cell culture models are increasingly utilized in cancer research to better replicate in vivo tumor microenvironments. This study examines the effects of different 3D scaffolding materials, including Matrigel, GelTrex, and the plant-based GrowDex, on prostate cancer cell lines, with a particular emphasis on neuroendocrine prostate cancer (NEPC). Four cell lines (LNCaP, LASCPC-01, PC-3, and KUCaP13) were cultured in these scaffolds to evaluate spheroid formation, cell viability, and gene expression. The results revealed that while all scaffolds supported cell viability, spheroid formation varied significantly: Matrigel promoted the most robust spheroids, especially for LASCPC-01, whereas GrowDex exhibited limitations for certain cell lines. Gene expression analysis indicated a consistent reduction in androgen receptor (AR) expression in LNCaP cells across all scaffolds, suggesting a potential shift towards a neuroendocrine phenotype. However, the expression of neuroendocrine markers varied depending on the scaffold and culture method, with the mini-domes method in Matrigel leading to decreased expression of both castration-resistant prostate cancer (CRPC) and NEPC markers. These findings highlight the scaffold-dependent variability in 3D culture outcomes and emphasize the need for standardized methodologies to ensure consistency and relevance in prostate cancer research.

**Highlights:**

- Comparing of Matrigel, GelTrex, and GrowDex in 3D prostate cancer culture
- Identifying scaffold-dependent spheroid formation and gene expression shifts
- Observing consistent AR reduction and variable neuroendocrine marker changes
- Highlighting the need for standard 3D culture methods in cancer studies

## 1. Introduction

Conventional 2D monolayer cell cultures have been an indispensable tool in cancer research due to their simplicity, fast growth, and high-throughput compatibility. However, 2D cultures cannot faithfully mimic the complex cell–cell and cell–extracellular matrix (ECM) interactions present in real tumors [1]. The absence of these three-dimensional interactions in 2D systems can alter cell differentiation, proliferation, gene and protein expression, and responses to stimuli [1]. Consequently, 2D cultures often fail to predict in vivo drug responses and tumor behavior. In contrast, 3D culture models provide a tissue-like architecture and microenvironment that bridge the gap between traditional cell culture and living tissues. By allowing cancer cells to grow as multicellular spheroids or organoids embedded in a supportive matrix, 3D models re-establish critical signaling pathways and biomechanical cues that govern tumor physiology [2]. These models have shown improved translational relevance, including drug response profiles more similar to tumor xenografts and patient tumors [3].

Prostate cancer research has increasingly adopted 3D models to study disease subsets and treatment responses. For example, Edmondson *et al.* demonstrated that LNCaP prostate cancer cells grown in a Matrigel-based 3D organoid form were significantly less sensitive to the chemotherapy drug docetaxel compared to 2D cultures [4]. Similarly, VanDeusen *et al.* developed a 3D model of NEPC by culturing NCI-H660 cells in a peptide hydrogel, and found that a higher dose of a RET inhibitor (AD80) was required to achieve 50% cell death in 3D vs. 2D [5]. These studies illustrate that 3D culture can alter therapeutic responsiveness, underscoring the value of 3D models for different prostate cancer subtypes. In fact, strong correlations have been observed between drug responses in 3D prostate culture models and tumor xenografts, suggesting 3D *in vitro* systems more accurately reflect *in vivo* tumor behavior than 2D monolayers [6].

A critical component of 3D culture systems is the scaffold or matrix that supports 3D growth. The current “gold standard” is Matrigel, a basement membrane extract from EHS mouse sarcoma rich in ECM proteins (approximately 60% laminin, 30% collagen IV, plus entactin and heparan sulfate proteoglycans) and growth factors [7]. Matrigel polymerizes at physiological temperature to form a gel, providing structural support and biochemical cues that promote cell proliferation and differentiation [8]. Geltrex is a similar ECM protein mixture with reduced growth factor content and lot-to-lot variability, making it a more standardized variant of Matrigel [9]. These matrices have enabled countless 3D culture applications including spheroid/organoid growth and drug delivery studies. However, reliance on Matrigel and other animal-derived matrices raises concerns about sustainability, reproducibility, and undefined components. Matrigel production requires substantial use of laboratory animals, and its complex composition (including unknown levels of growth factors and cytokines) can introduce experimental variability and confound results [10]. To address these issues, synthetic and plant-based hydrogels are being explored as alternative 3D scaffolds. One promising example is GrowDex, a hydrogel derived from nanofibrillar cellulose of birch trees [11]. GrowDex contains no exogenous proteins or growth factors, offering a biologically inert but tunable matrix that mimics the physical architecture of the ECM. Early studies have shown that GrowDex can support 3D cultures of various cell types with results comparable to Matrigel. For instance, smooth muscle and endothelial cells cultured in GrowDex formed structures with morphology and viability similar to those in Matrigel [12]. GrowDex’s consistency and defined composition (minimal batch variability) make it attractive for standardized 3D assays. Nevertheless, its efficacy with cancer cells, especially those requiring growth factor cues, needs further evaluation.

Neuroendocrine prostate cancer (NEPC) is a clinically important context and suitable to apply advanced 3D models. Prostate cancer is initially driven by androgens and is commonly treated with androgen deprivation therapy (ADT) and AR pathway inhibitors [13]. While these therapies are effective initially, tumors often develop resistance and progress to castration-resistant prostate cancer (CRPC). Most CRPCs remain AR-driven and respond to second-generation antiandrogens (enzalutamide, apalutamide, darolutamide) or androgen synthesis blockers (abiraterone). However, a subset of CRPC (∼15–20%) eventually loses dependency on AR signaling [14]. This shift can occur via a process of lineage plasticity or transdifferentiation wherein prostate adenocarcinoma cells adopt a neuroendocrine-like phenotype as a mechanism of treatment escape [15]. The result is therapy-induced NEPC (also called treatment-emergent small cell carcinoma), an aggressive AR-negative variant characterized by poor differentiation, rapid metastasis, and resistance to conventional AR-targeted therapies [16]. Clinically, NEPC is identified by low or absent AR and prostate-specific antigen (PSA) expression, coupled with the expression of neuroendocrine markers such as chromogranin A (CHGA), neuron-specific enolase (ENO2), neural cell adhesion molecule (NCAM1), and synaptophysin (SYP). NEPC commonly involves elevated expression of developmental transcription factors (e.g., SOX2, ASCL1) that drive the alternative lineage program [17] [18] [19]. This phenotype is associated with aggressive behavior and poor prognosis. It is estimated that among CRPC patients treated with potent AR inhibitors, around 17% eventually develop clinically apparent NEPC by histology, though some estimates range up to ∼25–30% when including mixed phenotypes [16]. Despite its importance, NEPC remains incompletely understood and lacks effective targeted therapies beyond platinum-based chemotherapy regimens [20].

Because NEPC typically arises under the pressure of AR pathway therapies, there is intense interest in understanding how the tumor microenvironment and cellular interactions contribute to this transdifferentiation. Recent studies suggest that 3D growth conditions might facilitate or reveal NEPC features that are not seen in 2D culture. The close cell–cell and cell–ECM contacts in spheroids can activate pathways involved in stemness, invasion, and therapy resistance – for example, Wnt signaling has been shown to be upregulated in 3D cultures and linked to more invasive phenotypes [21]. In glioma, 3D collagen scaffolds induced upregulation of genes related to stemness and migration compared to 2D, and similar mechanisms may operate in prostate cancer [22]. Indeed, prior work reported that LNCaP prostate cancer cells grown in 3D showed reduced AR protein and activation of noncanonical Wnt signals associated with antiandrogen resistance [23]. Moreover, ECM properties such as stiffness can influence lineage plasticity: a stiffer matrix can promote pathways that enable NE differentiation and castration resistance [24]. These insights led us to hypothesize that 3D culture might accentuate or induce NEPC-like characteristics in prostate cancer cells, thereby serving as a useful model to study NEPC development in vitro.

In this study, we systematically examine how different 3D culture matrices and methods affect prostate cancer cell phenotype, with a focus on NEPC-related traits. We cultured a panel of prostate cancer cell lines – LNCaP (AR+/NE– adenocarcinoma), 22Rv1 (AR-splice variant+/NE+ CRPC), LASCPC-01 (AR–/NE+ NEPC), PC-3 (AR– adenocarcinoma with some NE features), and KUCaP13 (AR–/NE+ NEPC) – under various 3D conditions. Three scaffold materials were compared: Matrigel and Geltrex (animal-derived ECM gels) versus GrowDex (plant-based cellulose hydrogel). We also compared two 3D culture techniques: a sandwich method (cells embedded between layers of hydrogel in multiwell plates) and a mini-dome method (small Matrigel droplets or “domes” seeded in dishes, per organoid culture protocols). We assessed outcomes including spheroid formation efficiency, cell viability and yield, and expression of key marker genes and proteins (AR, PSA for luminal/CRPC phenotype; CHGA, ENO2, NCAM1, SYP, ASCL1, STEAP1 for NEPC). Our goal was to determine how scaffold composition and culture methodology influence the short-term phenotype of prostate cancer cells, particularly regarding any shift toward or away from NEPC characteristics. These findings provide guidance for selecting 3D culture systems in prostate cancer research and highlight the need for standardized 3D protocols to reliably model disease progression and treatment resistance.

## 2. Materials and Methods

### 2.1. Cell lines and culture conditions

We used five human prostate cancer cell lines representing different disease states and phenotypes: LNCaP, 22Rv1, PC-3, LASCPC-01, and KUCaP13. The characteristics of each cell line are summarized in **Table 1**. LNCaP (AR-positive, PSA-secreting) and 22Rv1 (AR-positive, including the constitutively active AR-V7 splice variant) are models of prostate adenocarcinoma and castration-resistant prostate cancer (CRPC). PC-3 is an AR-negative adenocarcinoma line that exhibits some neuroendocrine marker expression (e.g., chromogranin A and ENO2). LASCPC-01 and KUCaP13 are AR-negative, PSA-negative cell lines derived from neuroendocrine prostate cancer (NEPC) tumors, thus serving as models of the neuroendocrine phenotype. All cell lines were cultured in both traditional 2D and various 3D conditions as described below. For routine maintenance and 2D experiments, LNCaP, 22Rv1, PC-3, and KUCaP13 were grown in RPMI-1640 medium supplemented with 10% fetal bovine serum (FBS) and 1% penicillin–streptomycin. LASCPC-01 was maintained in HITES medium (RPMI-1640 base with 5 μg/mL insulin, 10 μg/mL transferrin, 30 nM sodium selenite, 10 nM hydrocortisone, 10 nM β-estradiol, and 10 mM HEPES) to support its growth factor requirements. All cultures were kept in a humidified incubator at 37 °C with 5% CO_2_.

**Table 1.** Prostate cancer cell lines used and their phenotypic features.

| Cell Line | Disease/Phenotype | AR/PSA status | NE marker status |
| --- | --- | --- | --- |
| **LNCaP** | Androgen-sensitive prostate adenocarcinoma (lymph node metastasis) | AR+, PSA+ | Low/negative NE markers |
| **PC-3** | Castration-resistant prostate adenocarcinoma (bone metastasis) | AR–, PSA– | Moderate NE markers |
| **22Rv1** | Castration-resistant prostate cancer (with AR-V7 variant) | AR+, PSA+ | Moderate NE markers |
| **LASCPC-01** | Treatment-induced NEPC (luminal-to-NE transdifferentiation model) | AR–, PSA– | Strong NE phenotype |
| **KUCaP13** | Neuroendocrine prostate cancer<br>(patient-derived) | AR–, PSA– | Strong NE phenotype |

### 2.2. 3D scaffold materials

Three biologically distinct scaffold materials were employed for 3D cultures: (1) Matrigel Growth Factor Reduced (GFR) Basement Membrane Matrix: a solubilized basement membrane extract from Engelbreth-Holm-Swarm (EHS) mouse sarcoma, containing predominantly laminin, collagen IV, entactin, and heparan sulfate proteoglycans. The GFR version has reduced levels of exogenous growth factors and cytokines, but still provides an ECM-rich environment. Matrigel was obtained from Corning and used according to manufacturer guidelines [25]. (2) Geltrex LDEV-Free Reduced Growth Factor Basement Membrane Matrix: a product similar to Matrigel (also derived from EHS mouse tumor) with a more defined composition and lower batch-to-batch variability in protein and growth factor content. Geltrex (Thermo Fisher Scientific) was handled and diluted in the same manner as Matrigel [25]. (3) GrowDex Hydrogel: a bioinert matrix of nanofibrillar cellulose derived from birch wood. GrowDex contains no mammalian proteins or growth factors, offering high lot-to-lot consistency. It mimics the physical aspects of the ECM and can be formulated at different concentrations to adjust stiffness. GrowDex was procured from UPM Biomedicals and prepared as per the supplier’s instructions [26]. For the sandwich culture method (used with Matrigel and Geltrex), we diluted the matrix stocks 1:2 with the appropriate culture medium to achieve a working protein concentration of ∼5 mg/mL, as recommended. GrowDex was used at two concentrations (0.5% and 0.25% w/v) in preliminary tests, but 0.5% was adopted as the standard concentration unless otherwise noted.

### 2.3. 3D culture methods

We explored two different 3D culture methodologies: a sandwich embed method for all three scaffolds, and a mini-dome (droplet) method for Matrigel only. All 3D cultures were initiated with single-cell suspensions and grown for short-term periods (5–7 days) prior to analysis.

**(1) 2D seeding for controls:** For each experiment, control 2D cultures were established in parallel. Adherent lines (LNCaP, 22Rv1, PC-3) were trypsinized to single cells; suspension-grown lines (LASCPC-01, KUCaP13) were collected directly. Cells were pelleted and resuspended in growth media. We seeded 2D controls at comparable densities to the 3D cultures (e.g., 1×10^6^ cells in a 10 cm dish for sandwich method experiments, or 2.5×10^5^ cells in one well of a 6-well plate for mini-dome experiments). Adherent cultures were maintained in standard plates (10 cm or 6-well format) with media changes every other day until harvest.
**(2) Sandwich method (Matrigel/Geltrex 3D cultures):** 3D sandwich cultures in Matrigel and Geltrex were prepared following Corning’s protocol for 3D ECM culture [25]. Briefly, 100 μL of cold diluted Matrigel or Geltrex solution was added to each well of a 48-well plate and allowed to solidify for 30 minutes at 37 °C, forming a base gel layer. Cells were mixed with additional matrix solution (e.g., 90 μL matrix + 10 μL cell suspension containing 100,000 cells) on ice, then 100 μL of this cell–matrix mixture was overlaid on the base layer and gelled at 37 °C for ∼45 minutes. After gelation, 100 μL of culture medium was gently added atop the second layer to nourish the embedded cells. Cultures were maintained at 37 °C, 5% CO_2_, with medium changes every 2 days, and harvested on Day 5 for viability and RNA analysis (and on Day 7 for any protein analysis). The same procedure was used for both Matrigel and Geltrex.
**(3) GrowDex 3D Culture:** GrowDex cultures were set up according to the manufacturer’s guidelines [26]. We mixed cells with sterile GrowDex to a final volume of 300 μL (containing 0.5% GrowDex and ∼1×10^5^ cells). This mixture was aliquoted into 3 wells of a 48-well plate (∼100 μL per well) to form a uniform 3D cell suspension. An additional 100 μL of culture medium was carefully layered on top of each well to supply nutrients. GrowDex cultures were fed by replacing the top medium layer every 2 days. Cells were grown for 5 days before harvest.
**(4) Mini-dome 3D culture (Matrigel droplet method):** To test a different 3D seeding approach, we employed an organoid-style dome culture in Matrigel (adapted from ATCC Organoid Culture Guide) [27]. After trypsinization or collection, 250,000 cells were pelleted and resuspended in 100 μL of cold Matrigel per sample. Using a chilled pipette, ten 10-μL droplets (“mini-domes”) of the cell–Matrigel suspension were placed onto the surface of a pre-warmed 6-well plate (i.e. ∼10,000 cells per dome). The plate was then inverted and incubated for 15–30 minutes to allow the domes to solidify, anchoring the 3D microtissues to the plate. After gelation, 2 mL of warm culture medium was gently added to each well, fully covering the domes without dislodging them. Medium was changed every other day. These 3D cultures were harvested on Day 7 for analysis (slightly longer culture than the sandwich method, to allow sufficient spheroid growth in domes).
**(5) Cell imaging:** To monitor spheroid formation and morphology, we imaged cultures on Day 5 after seeding. Bright-field images were taken for each condition using a Leica inverted microscope (10× objective). LNCaP and LASCPC-01 spheroids in Matrigel, Geltrex, and GrowDex were photographed to compare their size and density. For the mini-dome experiment, serial images were taken over the 7-day period to observe spheroid development in Matrigel domes vs. 2D. All images used consistent exposure and magnification; scale bars represent 25–50 μm as indicated.

### 2.4. Harvesting cells from 3D cultures

Recovering viable cells from 3D matrices required different dissociation techniques tailored to each scaffold:

**(1) Matrigel/Geltrex (sandwich cultures):** We used enzyme-mediated digestion with dispase in Hanks’ Balanced Salt Solution (HBSS). Specifically, 150 μL of a 1× dispase solution (5 U/mL; diluted in HBSS) was added to each well of the 48-well plate after removing the top medium. The plate was incubated at 37 °C for 1 hour to partially dissolve the ECM gel. Then, the contents of each well (cells + partially digested matrix) were gently pipetted up and down, transferred to a tube, and centrifuged at 300×g for 5 minutes at 4 °C. The cell pellet was washed once with cold PBS and centrifuged again. Finally, the pellet was resuspended in fresh media for downstream assays. This protocol efficiently released cells from the soft Matrigel and Geltrex gels.
**(2) GrowDex:** We followed the manufacturer’s instructions using GrowDase, a cellulase enzyme that specifically degrades the cellulose hydrogel [28]. For each well (initially ∼100 μL of 0.5% GrowDex), we added an appropriate volume of GrowDase solution (at 1.5 U of enzyme per mg of GrowDex) in culture medium to match the well volume. The plate was incubated at 37 °C for at least 8 hours (or overnight), allowing the enzyme to completely digest the cellulose matrix. Subsequently, the contents were collected and spun at 300×g for 5 minutes (4 °C) to pellet the cells. The pellet was washed with PBS and centrifuged again to remove residual hydrogel and enzyme. Cells were then resuspended in medium for counting or RNA extraction.
**(3) 2D cultures:** Adherent 2D cultures (LNCaP, 22Rv1, PC-3) were harvested by gently scraping with a cell lifter (to avoid excessive trypsinization that could affect gene expression). Suspension 2D cultures (LASCPC-01, KUCaP13) were simply collected. All 2D samples were centrifuged (300×g, 5 min, 4 °C) and washed with PBS in the same manner as 3D samples. This ensured that 2D and 3D cells underwent similar handling before analysis.
**(4) Matrigel mini-domes:** We applied a mechanical dissociation approach to recover cells from Matrigel domes (since enzymatic methods risk cell loss given the small dome size). After removing culture medium, each well was gently scraped with a cell lifter to loosen the domes. We added cold PBS and used pre-wet pipette tips to carefully transfer the disrupted domes into 15 mL conical tubes. The domes were further broken up by pipetting and diluted with additional PBS (up to 15 mL). The tube was then centrifuged at 300×g for 5 minutes. The supernatant was aspirated, and the cell pellet was washed with PBS and pelleted again. The resulting pellet was resuspended in medium for viability counting or lysis for molecular analysis. This gentle mechanical method (adapted from organoid handling protocols) minimized cell damage while retrieving cells from the mini-domes. After dissociation, an aliquot of each sample was taken for cell count and viability assessment using trypan blue exclusion. A 1:1 mixture of cell suspension and 0.4% trypan blue was loaded into an automated cell counter slide. Live (unstained) and dead (blue-stained) cells were counted with a Thermo Countess II, and total viable cell numbers and % viability were recorded for each condition. The remaining cells were then processed for RNA or protein analyses as described below.

### 2.5. Quantitative Reverse Transcription PCR (qRT-PCR)

To quantify gene expression changes, we isolated RNA from 2D and 3D cultured cells and performed qRT-PCR for selected genes. Total RNA was extracted using a commercial spin column kit (BioBasic) following the manufacturer’s protocol [29]. The yield and purity of RNA were confirmed by spectrophotometry. Complementary DNA (cDNA) was synthesized from 500 ng – 1 μg of each RNA sample using a high-efficiency reverse transcription kit (All-In-One 5× RT MasterMix, Applied Biological Materials). Reactions were set up according to the supplier’s instructions [30]. Next, qPCR was conducted using SYBR Green chemistry on a Bio-Rad CFX96 system. Each 20 μL reaction contained 2 μL of diluted cDNA, 10 μL of 2× SYBR Green qPCR master mix, and 0.5 μM of forward and reverse primers for the target gene. Cycling conditions were: 95 °C for 2 min initial denaturation, followed by 40 cycles of 95 °C for 15 s and 60 °C for 60 s. A melt curve analysis confirmed specific amplification. We analyzed the expression of luminal/CRPC markers (AR, PSA, STEAP1) and NEPC markers (CHGA, ENO2, NCAM1, SYP, ASCL1) in the various cultures. GAPDH was used as the housekeeping gene for normalization. Primer sequences are provided in **Table 2**. Quantification of relative gene expression was done using the comparative ΔΔCq method [31]. All gene expression levels in 3D are presented as fold-change relative to the 2D condition. Each experiment was performed with at least two biological replicates, and qPCR for each gene was run in triplicate wells. We considered a gene significantly differentially expressed if the fold-change was >1.5 in either direction with a p-value ≤ 0.05.

**Table 2.**
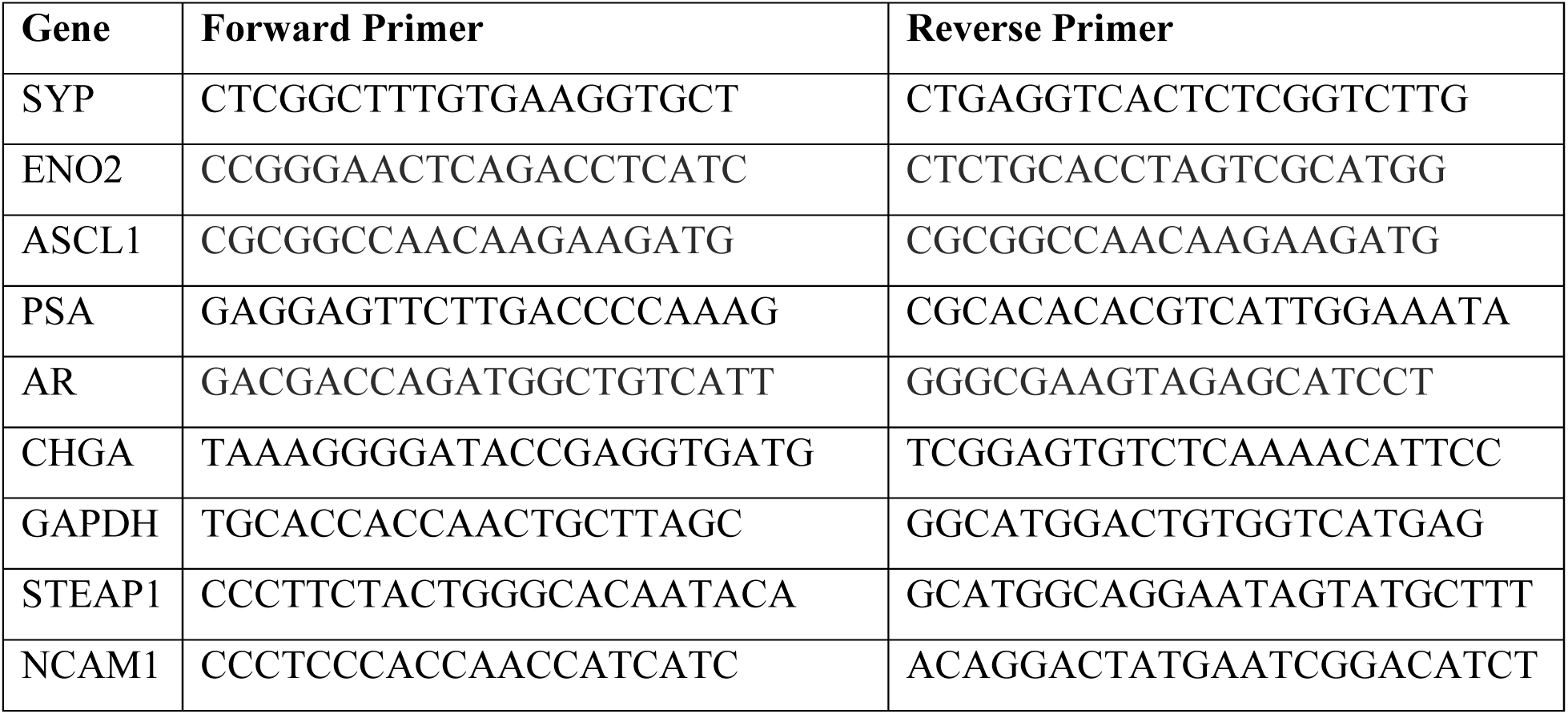
qRT-PCR primer sequences for examined marker genes.

### 2.6. Western blotting

Cell pellets were lysed in ice-cold RIPA buffer supplemented with protease and phosphatase inhibitors. Lysates were briefly sonicated (5 s) to shear DNA and reduce viscosity. After a 30 min incubation on ice, lysates were clarified by centrifugation (10 min at 14,000×g). Protein concentration was determined using a BCA protein assay kit (Thermo Fisher) [32]. Equal amounts of protein (20–30 μg) from each sample were mixed with Laemmli buffer, boiled, and loaded onto a 10% SDS-PAGE gel. Following electrophoresis, proteins were transferred to a PVDF membrane using a semi-dry transfer apparatus. Membranes were blocked in 5% nonfat milk in TBS-Tween for 1 hour at room temperature. Blots were probed overnight at 4 °C with primary antibodies against AR (Cell Signaling Technology #5153) and PSA (Cell Signaling Technology #5365). Histone H3 (Abcam ab21054) or GAPDH (Cell Signaling Technology #2118) were used as loading controls. After washing, membranes were incubated with HRP-conjugated secondary antibodies, followed by adding chemiluminescent substrate and detecting signals with the ChemiDoc imaging system (Bio-Rad).

### 2.7. Statistical Analysis

All quantitative data are presented as mean ± standard deviation. Experiments were repeated at least twice independently. Statistical significance between groups was determined using appropriate tests in GraphPad Prism. For comparisons of multiple 3D conditions to 2D (e.g., gene expression across three matrices vs 2D), we used one-way or two-way ANOVA with post hoc tests as indicated in figure legends. For direct pairwise comparisons (e.g., 2D vs 3D mini-dome), unpaired two-tailed t-tests with Welch’s correction were applied. A p-value < 0.05 was considered statistically significant.

## 3. Results

### 3.1. Scaffold composition influences spheroid formation and cell recovery in 3D culture

We first examined the ability of each matrix (Matrigel, Geltrex, GrowDex) to support spheroid growth of prostate cancer cells using the 3D sandwich method. LNCaP cells readily formed multicellular spheroids in all three scaffold types (**Fig. 1A**). Matrigel produced the largest and most numerous LNCaP spheroids, followed by Geltrex (slightly smaller or fewer spheroids). GrowDex also supported robust LNCaP spheroid formation, though spheroids appeared somewhat sparser in number compared to the protein-based matrices. Notably, the cellulose fibers in GrowDex created a background that made it challenging to visualize all spheroids.

**Figure 1.**
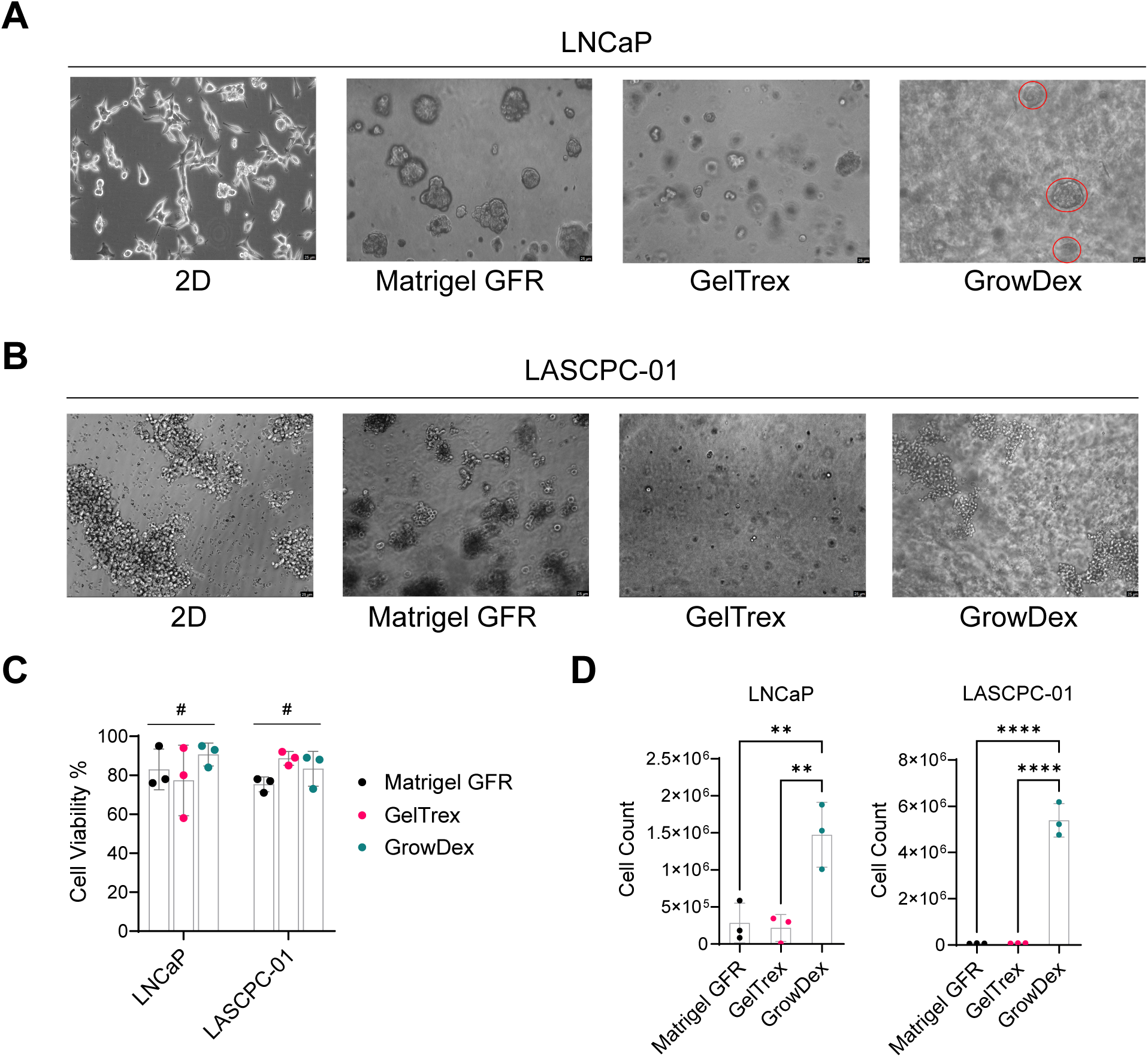
Spheroid formation and cell recovery in different 3D scaffold materials for LNCaP and LASCPC-01. **(A)** Morphology of LNCaP in 2D culture and 3D culture (Matrigel, Geltrex, GrowDex). Scale bars = 25 µm. **(B)** Morphology of LASCPC-01 in 2D culture and 3D culture (Matrigel, Geltrex, GrowDex). Scale bars = 25 µm. **(C)** Cell viability of LNCaP and LASCPC-01 on Day 5 in each 3D condition. ^#^ p > 0.05, two-way ANOVA. **(D)** Total cell counts of LNCaP and LASCPC-01 from different 3D matrices. **p < 0.01, ****p <0.0001, one-way ANOVA with Turkey’s multiple comparison test.

In contrast, LASCPC-01 cells showed more limited spheroid formation in 3D (**Fig. 1B**). In 2D culture, LASCPC-01 formed floating clumps rather than a monolayer. In 3D Matrigel, we observed some spheroid-like structures by Day 5, although they were generally smaller and fewer than the ones formed by LNCaP. Geltrex failed to induce true spheroids in LASCPC-01; the cells tended to form irregular aggregates or loose clusters, but not the rounded spheroids seen in Matrigel. Similarly, in GrowDex, LASCPC-01 aggregates resembled the floating clusters from 2D, with no improvement in cohesion or spheroid architecture. These results suggest that exogenous growth factors or matrix cues are necessary for LASCPC-01 to form spheroids.

We next assessed cell viability and recovery from the different 3D cultures on Day 5. Trypan blue exclusion showed that cell viability remained high (75–90%) in all conditions for both LNCaP and LASCPC-01 (**Fig. 1C**), with no significant differences between Matrigel, Geltrex, or GrowDex. Thus, the various matrices were comparably biocompatible, and neither the absence of protein in GrowDex nor the presence of mouse proteins in Matrigel/Geltrex overtly affected short-term cell survival. However, the number of cells recovered after matrix dissociation differed substantially by scaffold (**Fig. 1D**). For both LNCaP and LASCPC-01, the cell counts obtained from GrowDex cultures were significantly higher (by ∼2-fold) than those from Matrigel or Geltrex. This was initially surprising given that Matrigel had the highest spheroid density microscopically. We attribute this difference to the efficiency of cell harvest methods. The cellulase enzyme (GrowDase) completely solubilized the GrowDex matrix, likely releasing nearly all embedded cells. In contrast, the dispase-based recovery from Matrigel/Geltrex might not liberate every cell (some cells can remain trapped in undigested matrix clumps). Therefore, the higher yield from GrowDex underscores a practical advantage: the ease of fully recovering cells for downstream assays. It also suggests that optimizing dissociation protocols (e.g., stronger enzymes or longer digestion for ECM gels) could improve cell yields from Matrigel/Geltrex. Importantly, all conditions yielded sufficient cells for RNA and protein analyses, but GrowDex had a clear edge in harvest efficiency, which could be beneficial for scaling up 3D cultures or performing single-cell analyses.

### 3.2. Differential gene expression in 3D Cultures: AR suppression and neuroendocrine marker changes

We next analyzed how 3D culturing in different matrices affected the expression of key prostate cancer genes in LNCaP (adenocarcinoma) and LASCPC-01 (NEPC) cells. Using qRT-PCR, we measured the fold-change in mRNA levels for AR and its transcriptional target PSA (KLK3), as well as several neuroendocrine or plasticity-related markers, in Day 5 3D cultures relative to 2D controls. The results revealed both common trends and matrix-specific effects on gene expression. In LNCaP cells, one consistent finding was a strong reduction (by ∼60–80%) in AR mRNA in all 3D conditions compared to 2D (**Fig. 2A**). Correspondingly, PSA was also markedly downregulated in Matrigel and GrowDex (**Fig. 2B**). Consistent with the mRNA data, AR and PSA proteins in LNCaP cells were greatly diminished in 3D (Matrigel) relative to 2D (**Fig. 2C**). The neuroendocrine marker gene ENO2 (encoding neuron-specific enolase) was a mildly expressed gene in LNCaP, and it was found significantly upregulated in LNCaP spheroids in Geltrex and GrowDex (**Fig. 2D**). The two other neuroendocrine marker genes CHGA and SYP were absent in LNCaP and did not show meaningful changes in 3D culture (data not shown). Overall, LNCaP exhibited a shift toward an AR-low, NE marker-high state in 3D, especially pronounced in certain matrices.

**Figure 2.**
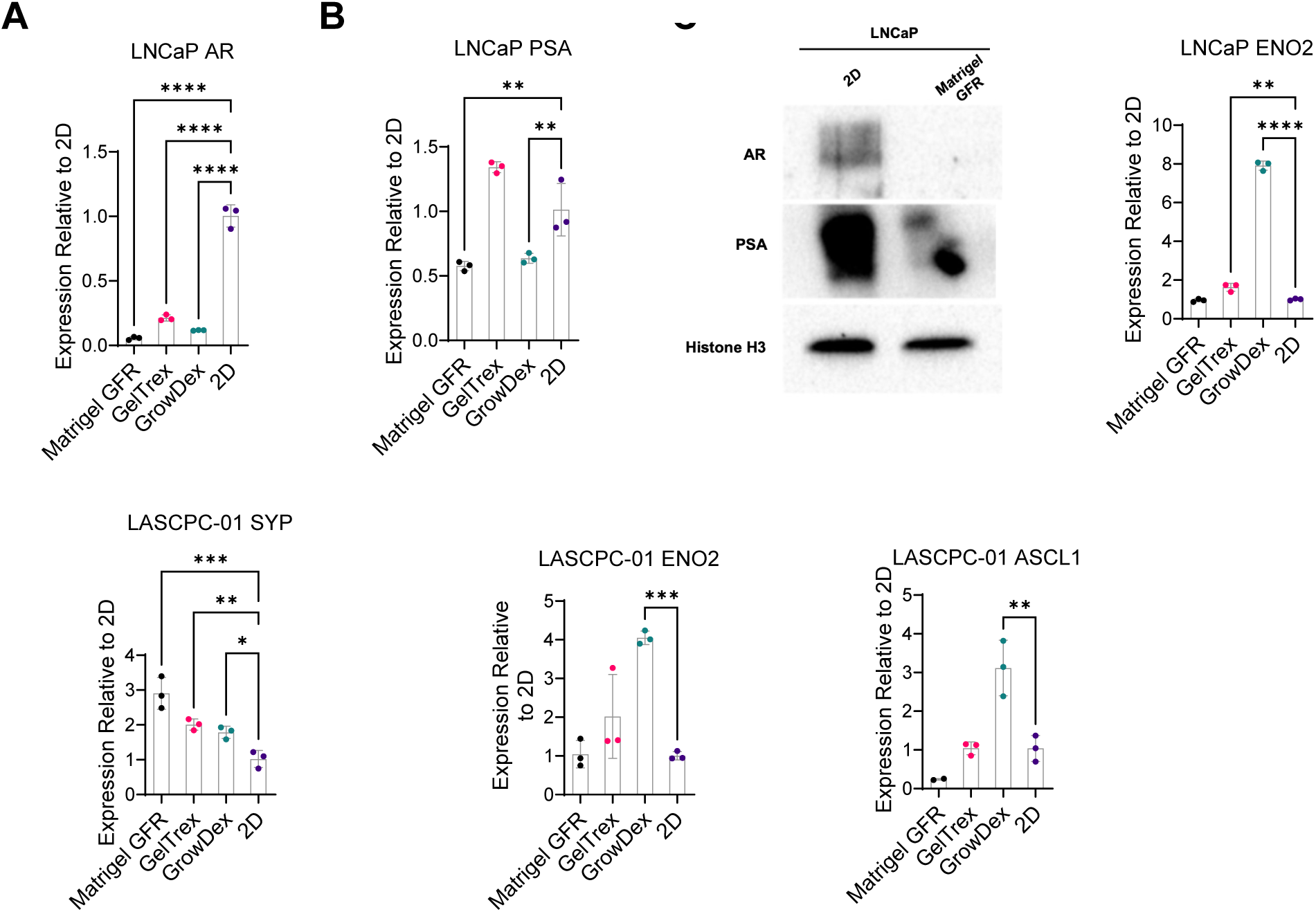
Gene and protein expression changes in LNCaP and LASCPC-01 in 3D culture. **(A, B)** Relative mRNA expression of AR and PSA in LNCaP after 5 days in 3D culture as relative to 2D control. **(C)** Western blot analysis of AR and PSA protein levels in LNCaP after 7 days in 3D Matrigel versus 2D culture. Histone H3 was used as a loading control. **(C)** Relative mRNA expression of ENO2 in LNCaP after 5 days in 3D culture as relative to 2D control. (E-G) Relative mRNA expression of AR and PSA in LASCPC-01after 5 days in 3D culture as relative to 2D control. **p < 0.01, ***p < 0.001, ****p <0.0001, one-way ANOVA with Dunnett’s multiple comparison test.

In LASCPC-01 (the AR-negative NEPC model), we focused on NE markers, since AR and PSA are not expressed. SYP mRNA was consistently elevated in 3D cultures of LASCPC-01 relative to 2D (**Fig. 2E**). In GrowDex, LASCPC-01 exhibited significant upregulation of both ENO2 and ASCL1 compared to 2D, whereas in Matrigel and Geltrex those genes remained unchanged (**Fig. 2F-G**). These findings emphasize that even for an established NEPC cell line, the 3D microenvironment can alter gene expression profiles. In summary, LASCPC-01 in 3D may enhance specific neuroendocrine markers (e.g. SYP), but the magnitude of change in certain neuroendocrine genes (ENO2, ASCL1) varied with the scaffold.

### 3.3. Induction of neuroendocrine markers by GrowDex in PC-3 and KUCaP13

Given the distinct behaviors seen in LNCaP versus LASCPC-01 in different matrices, we extended our analysis to the other two cell lines, PC-3 and KUCaP13, using the sandwich 3D method in Geltrex and GrowDex. We selected these two lines as a follow-up because PC-3 and KUCaP13 both lack AR, PC-3 is adherent and only partially NE, and KUCaP13 is a bona fide NEPC line. We wanted to see if these cells would mirror the trends observed above or exhibit unique responses. PC-3 in 2D grows as a strongly adherent, spindle-shaped cell layer (**Fig. 3A**). In Geltrex, PC-3 formed small spheroids or clusters with some residual attached cells forming web-like extensions (**Fig. 3A**), suggesting partial adhesion to the matrix or incomplete detachment into spheroids. In GrowDex, PC-3 formed diffuse, small aggregates scattered in the cellulose matrix (**Fig. 3A**). These aggregates were much less compact than LNCaP spheroids in GrowDex, indicating that PC-3’s spheroid-forming capacity is limited. Regarding gene expression, NEPC markers (SYP, ENO2, CHGA, and ASCL1) were significantly upregulated in GrowDex compared to 2D, and while they were also elevated in GelTrex, the increase was less pronounced (**Fig. 3B**). This suggests that aggregate formation may correlate with NEPC marker expression in PC-3 cells.

**Figure 3.**
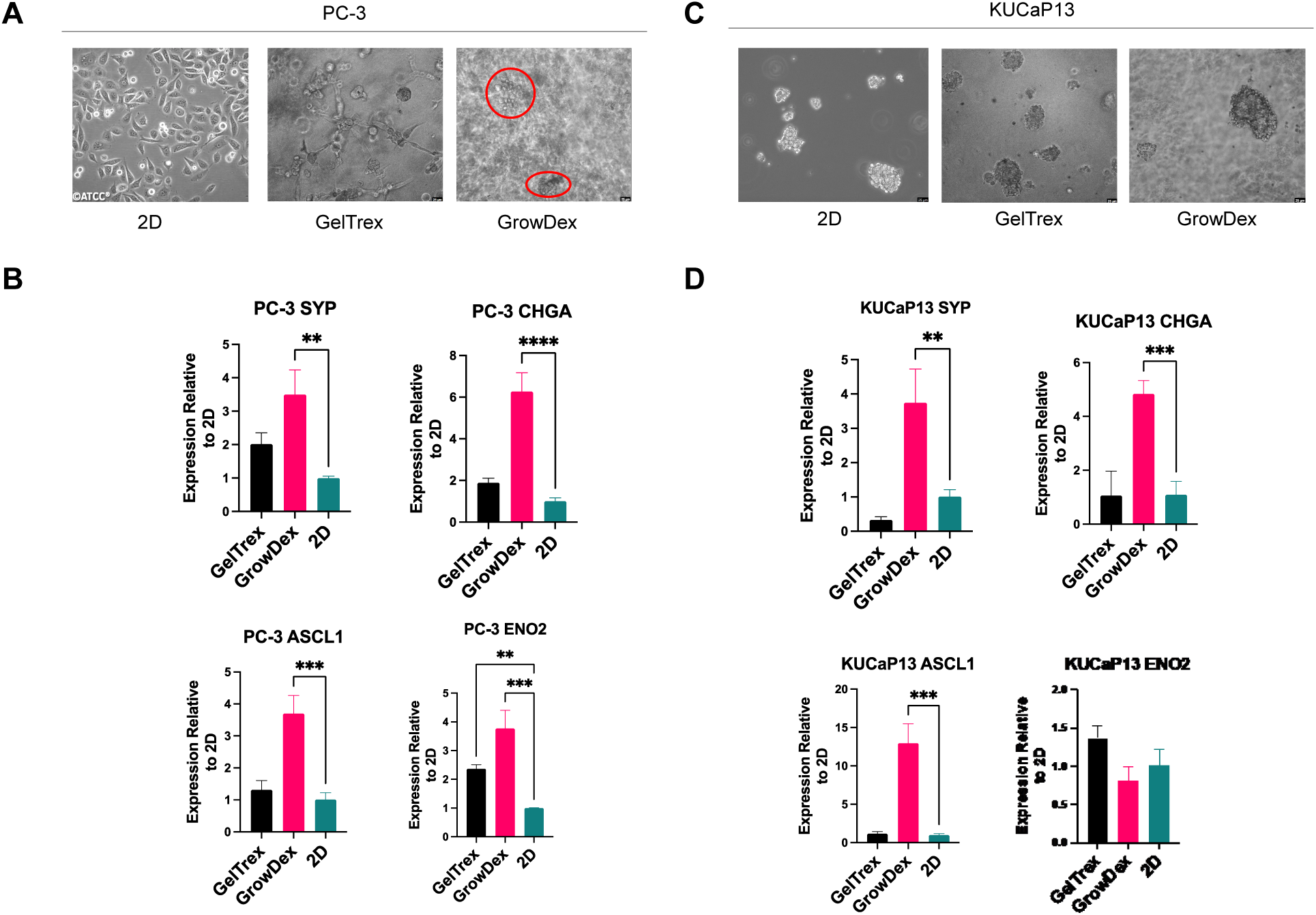
Morphology and neuroendocrine marker expression in PC-3 and KUCaP13 across different 3D scaffolds. **(A)** Morphology of PC-3 in 2D culture and 3D culture (Geltrex, GrowDex). Scale bars = 25 µm. **(B)** Relative mRNA expression of NEPC-related genes in PC-3 across different 3D conditions, normalized to 2D. **(C)** Morphology of KUCaP13 in 2D culture and 3D culture (Geltrex, GrowDex). Scale bars = 25 µm. **(D)** Relative mRNA expression of NEPC-related genes in KUCaP13 across different 3D conditions, normalized to 2D. **p < 0.01, ***p < 0.001, ****p <0.0001, one-way ANOVA with Dunnett’s multiple comparison test.

KUCaP13 in 2D grows as non-adherent dense clusters (similar to LASCPC-01) (**Fig. 3C**). When cultured in Geltrex or GrowDex, KUCaP13 also formed dense aggregates (**Fig. 3C**). Interestingly, the aggregates in Geltrex and GrowDex looked quite similar to each other and to the original 2D clusters, implying that KUCaP13’s growth pattern is inherently cluster-forming and not strongly altered by the matrix type. As observed in PC-3, NEPC markers (SYP, CHGA, and ASCL1) were significantly upregulated in GrowDex compared to both 2D and GelTrex (**Fig. 3D**). Meanwhile, ENO2 expression levels remained consistent across all conditions, indicating that this gene’s expression in KUCaP13 is not strongly affected by 3D culture. Collectively, these results indicate that 3D culture in GrowDex vs Geltrex can differentially modulate neuroendocrine features of prostate cancer cells that lack AR. Both PC-3 and KUCaP13 tend to ramp up NE-associated genes when grown in 3D, but a matrix with minimal biochemical signals (GrowDex) produced a more pronounced effect than an ECM protein matrix (Geltrex). It is possible that the presence of laminin/collagen and residual growth factors in Geltrex provides signals that keep cells slightly more “epithelial” or differentiated, whereas GrowDex’s inertness might impose stress that pushes cells toward a neuroendocrine-like gene program.

### 3.4. Matrigel “Mini-Dome” 3D culture globally suppresses CRPC and NEPC markers

Next, we investigated how an alternative 3D culturing technique – the Matrigel mini-dome method – would affect prostate cancer cells, in comparison to the sandwich Matrigel method above. In this experiment, we focused on LNCaP, 22Rv1, and LASCPC-01 to cover AR-positive and AR-negative scenarios. We cultured each cell line either in conventional 2D or in 3D Matrigel domes for 7 days. The spheroid morphology observed in LNCaP and LASCPC-01 cultures using the mini-dome method closely resembled that seen with the sandwich method (**Fig. 4A** vs **Fig. 1A**). In 2D culture, distinct proliferation patterns were evident among the cell lines, with LNCaP and 22Rv1 exhibiting adherent growth, whereas LASCPC-01 grew in suspension. However, in 3D culture, all cell lines formed relatively similar spheroid-like structures within the Matrigel scaffold. Notably, the LASCPC-01 spheroids appeared denser, consistent with its higher proliferation rate (doubling times: LNCaP, 60–72 hrs; 22Rv1, 35–40 hrs; LASCPC-01, 18 hrs). After 7 days, we dissociated the cultures, and the cell viabilities were slightly lower in 3D than in 2D (averaging ∼80% viable in 3D vs >90% in 2D for each line). This minor viability drop could result from the more rigorous recovery process from Matrigel, or from some cells in spheroids experiencing nutrient gradients. Nonetheless, viability remained sufficiently high for analysis.

**Figure 4.**
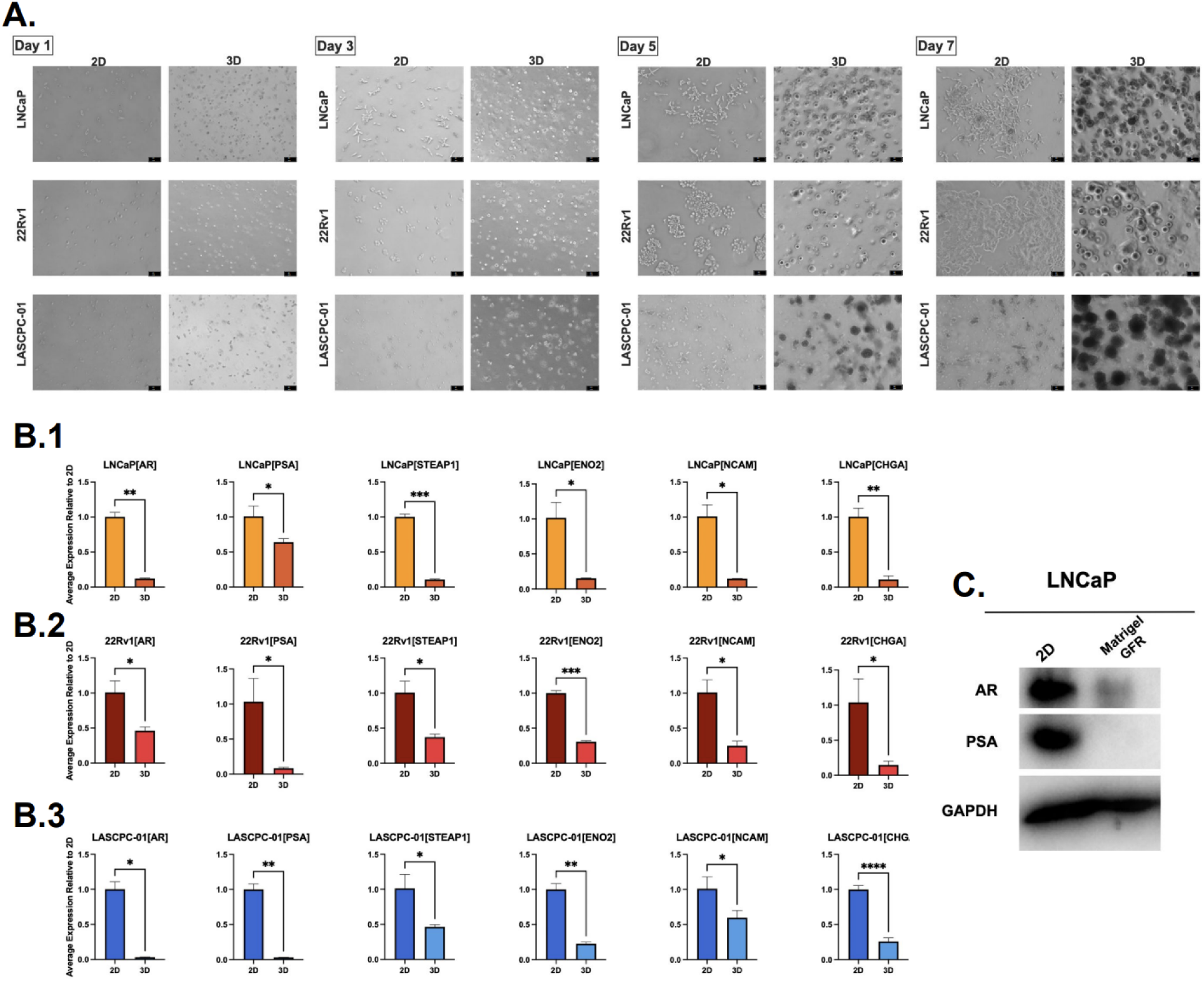
Morphology and gene marker expression changes in LNCaP, 22Rv1, LASCPC-01 in Matrigel mini-dome culture. **(A)** Bright-field images of LNCaP, 22Rv1, and LASCPC-01 in 2D and Matrigel mini-domes on different time points. Scale bars = 50 µm. **(B)** Relative mRNA expression of key prostate cancer markers in LNCaP (B.1), 22Rv1 (B.2), and LASCPC-01 (B.3) after 7 days in Matrigel mini-domes, normalized to 2D. Statistical significance determined by unpaired t-tests with Welch’s correction (*p < 0.05, **p < 0.01, ***p < 0.001, ****p < 0.0001). **(C)** Western blot analysis of AR and PSA protein levels in LNCaP after 7 days in Matrigel mini-domes versus 2D culture. GAPDH was used as a loading control.

The outcome of the mini-dome 3D culture was unexpectedly uniform across all cell lines: we observed a collective downregulation of both AR/CRPC markers and NEPC markers in 3D relative to 2D (**Fig. 4B**). Specifically, for each of LNCaP, 22Rv1, and LASCPC-01, the mRNA levels of AR, PSA, and STEAP1 (an AR-regulated gene often elevated in CRPC) were significantly lower in 3D (dropping to 10–30% of 2D levels). Simultaneously, the neuroendocrine-associated genes ENO2, NCAM1, and CHGA were also decreased in 3D compared to 2D (**Fig. 4B**). Thus, unlike the sandwich 3D results where NE markers often went up as AR went down, here both phenotypic categories (luminal/CRPC and NEPC markers) dropped in tandem under the mini-dome condition.

We corroborated the gene expression findings at the protein level for LNCaP. Western blot analysis of LNCaP from 2D vs 3D mini-domes showed that AR and PSA proteins were strongly decreased in 3D (**Fig. 4C**), consistent with the qRT-PCR results. These protein data confirm that the AR pathway is repressed in the mini-dome 3D environment, in line with the earlier observation from sandwich 3D.

## 4. Discussion

This study highlights how 3D culture systems significantly influence prostate cancer cell behavior, with outcomes dependent on both the scaffold type and culture method. We evaluated the feasibility of a plant-based hydrogel (GrowDex) as an alternative to animal-derived matrices and examined how different 3D conditions affect expression of AR and NEPC markers. Our findings underscore the need for careful selection and standardization of 3D models to improve their reliability in prostate cancer research.

GrowDex successfully supported spheroid formation in three of four prostate cancer cell lines (LNCaP, PC-3, and KUCaP13), yielding viable aggregates comparable to Matrigel and GelTrex. A key advantage was the ease of cell recovery using enzymatic digestion (GrowDase), which produced higher cell yields than Matrigel/GelTrex—beneficial for downstream assays requiring maximum recovery. Additionally, spheroids in GrowDex remained more discrete and size-controlled over time, contrasting with Matrigel, where rapid, uncontrolled spheroid growth led to fusion, a pattern previously observed in MCF-7 breast cancer cells [33]. The absence of growth factors in GrowDex likely contributed to its ability to sustain controlled spheroid formation. However, GrowDex was not universally suitable. LASCPC-01, an NEPC cell line, failed to form spheroids in GrowDex, likely due to its reliance on exogenous growth factors. Similarly, GelTrex, which has reduced growth factors compared to Matrigel, did not support spheroid formation for LASCPC-01, suggesting that some NEPC cells require additional biochemical cues. Functionalizing GrowDex with ECM proteins, such as fibronectin or vitronectin, has improved spheroid formation in fibroblasts and may enhance NEPC cell viability [34]. While GrowDex presents a sustainable alternative to Matrigel for many applications, further optimization is required for its broader use in prostate cancer research.

Our gene expression analysis revealed significant variability across 3D scaffolds. For example, LNCaP exhibited increased neuroendocrine marker (ENO2) expression in GrowDex and GelTrex but not in Matrigel, while LASCPC-01 showed differential expression across all matrices. These results suggest that 3D culture effects are not solely dictated by three-dimensionality but also by scaffold-specific biochemical and mechanical properties. This aligns with prior studies demonstrating that gene expression patterns in 3D cultures depend on factors such as ECM stiffness, degradability, and stress relaxation [35]. Given the inconsistencies observed, results from one 3D system may not be generalizable to another, reinforcing the need for detailed reporting of 3D culture conditions in publications.

A key observation was that 3D culture broadly suppressed AR expression, particularly in LNCaP, suggesting a shift toward a neuroendocrine-like phenotype. Previous studies have shown that 3D environments can promote lineage plasticity and resistance to therapy, as seen in glioma models where Wnt pathway activation was induced in 3D culture [22]. Similarly, prostate cancer cells in 3D scaffolds have exhibited reduced AR expression and increased Wnt signaling, a hallmark of treatment resistance [23]. In our study, NE marker upregulation was modest but consistent across some conditions, particularly in GrowDex. This suggests that 3D culture alone can activate pathways associated with lineage plasticity, potentially mimicking stress conditions seen during androgen deprivation therapy. However, the effect was not universal—some NE markers were not induced in LNCaP, and in the mini-dome method, NE markers were actually suppressed. Thus, 3D culture may predispose cells to neuroendocrine differentiation but is insufficient alone to drive full transdifferentiation without additional selective pressures such as antiandrogen treatment. Future studies could test whether treating 3D-cultured LNCaP spheroids with enzalutamide accelerates NE marker induction, creating a more physiologically relevant model of therapy-induced NEPC.

Interestingly, established NEPC cell lines (LASCPC-01 and KUCaP13) did not uniformly enhance their NE phenotype in 3D culture. Moreover, our mini-dome method led to a paradoxical suppression of NEPC markers, possibly due to factors such as Matrigel’s cytokine content, cell crowding, or nutrient saturation. Matrigel is rich in ligands like TGF-β and EGF, which may influence NE marker expression. These findings illustrate that even within the same 3D framework, small methodological variations can yield divergent outcomes, emphasizing the need for experimental standardization. The discrepancies between the sandwich and mini-dome methods further highlight the complexities of 3D culture. While both methods produced similar spheroid morphology, gene expression profiles differed significantly. This variability likely stems from differences in matrix geometry, oxygen diffusion, seeding density, and matrix retention of growth factors. The field lacks standardized 3D culture protocols, with widespread use of diverse methods such as hanging drops, hydrogels, microfluidics, and bioreactors, each introducing additional variables. Our results suggest that if 3D culture is to be widely adopted for prostate cancer research, a concerted effort toward protocol standardization is necessary.

These findings have implications for drug testing in 3D models. Given that AR expression was consistently lower in 3D cultures, AR-targeted therapies like enzalutamide may appear less effective in these models due to reduced target availability. Conversely, NEPC marker changes could influence sensitivity to other treatments. The unexpected suppression of both CRPC and NEPC markers in mini-domes raises questions about what phenotype these cells adopt in that setting—potentially a quiescent or indeterminate state. Testing drug responses in such a condition would be informative, as it may resemble a transition state between adenocarcinoma and NEPC.

In conclusion, our study underscores that 3D culture is not a monolithic technique, and scaffold selection, culture method, and biochemical environment all influence prostate cancer cell behavior. GrowDex presents a promising, sustainable alternative for 3D culture but requires further optimization for certain NEPC models. The broad reduction in AR expression across 3D conditions suggests that these models may better reflect certain aspects of advanced prostate cancer, including therapy resistance. However, methodological inconsistencies pose challenges, and without standardization, 3D cultures may yield conflicting or misleading data. Future studies should prioritize systematic comparisons of 3D methods and scaffolds to establish best practices for modeling prostate cancer progression and treatment response. By refining these approaches, 3D culture systems can become more effective tools for understanding lineage plasticity and developing targeted therapies for advanced prostate cancer.

## Acknowledgments

We thank the Lu lab members for their essential comments and suggestions during this work. This work was supported by Glynn Family Honors Program (E.F.), Interdisciplinary Interface Training Program (Z.Z.), and Boler Family Foundation (X.L.) at University of Notre Dame. Other support included National Institutes of Health grants R01CA248033 and R01CA280097 (X.L.), and Department of Defense grants W81XWH2010312, HT94252310010 and HT94252310613 (X.L.).

